# Out of Africa by spontaneous migration waves

**DOI:** 10.1101/378695

**Authors:** Paul D. Bons, Catherine C. Bauer, Hervé Bocherens, Tamara de Riese, Dorothée G. Drucker, Michael Francken, Lumila Menéndez, Alexandra Uhl, Boudewijn P. van Milligen, Christoph Wiβing

## Abstract

Hominin evolution is characterized by progressive regional differentiation, as well as migration waves, leading to anatomically modern humans that are assumed to have emerged in Africa and spread over the whole world. Why or whether Africa was the source region of modern humans and what caused their spread remains subject of ongoing debate. We present a spatially explicit, stochastic numerical model that includes ongoing mutations, demic diffusion, assortative mating and migration waves. Diffusion and assortative mating alone result in a structured population with relatively homogeneous regions bound by sharp clines. The addition of migration waves results in a power-law distribution of wave areas: for every large wave, many more small waves are expected to occur. This suggests that one or more out-of-Africa migrations would probably have been accompanied by numerous smaller migration waves across the world. The migration waves are considered “spontaneous”, as the current model excludes environmental or other factors. Large waves preferentially emanate from the central areas of large, compact inhabited areas. During the Pleistocene, Africa was the largest such area most of the time, making Africa the statistically most likely origin of anatomically modern humans, without a need to invoke additional environmental or ecological drivers.

## Introduction

Hominins are generally supposed to have originated in Africa and settled most of Africa and the southern half of Eurasia in the early Pleistocene [1–5]. Fossil evidence suggests that earliest *Homo sapiens* appeared in Africa during the late Middle Pleistocene (Jebel Irhoud, Omo and Herto [6–8]). Anatomically modern humans (AMH) emerged, spread out of Africa during the Late Pleistocene and now occupy the whole world as the only *Homo* species [9–13]. In the intervening time, a number of *Homo* species, such as Neanderthals, Denisovans, *Homo erectus, H. heidelbergensis, H. ergaster*, etc. existed, developed and disappeared again [14–21]. Pleistocene human evolution is thus characterised by differentiation, speciation, migration waves and extinction events. Most authors assume that the out-of-Africa spread of AMH involved one or more migration waves that replaced *Homo* species that existed at the time with only limited genetic admixture, such as between Neanderthals and AMH [22–23]. Many studies have addressed the timing and origin of migration waves, as well as migration paths [e.g. 24,22,25,26,13]. Apart from the major out-of-Africa event(s), AMH populations experienced several more migration waves within already populated areas, such as Africa [27–28] and Europe [29–30]. Considering this, it is not unlikely that more migration waves occurred in the Pleistocene, but the sparse fossil record still makes it difficult to detect any.

Assuming that migration waves did happen, the question arises what caused them, in particular the spread of AMH. Most authors favour some competitive advantage of AMH over other *Homo* species [31–33]. With climate change now central in the scientific discourse, many recent studies suggest that climate played an important role in the environmental changes making AMH more competitive than other *Homo* species, or allowing opening ecological corridors for dispersal of *Homo sapiens* out of Africa [34–42]. An opposite view is that AMH and Neanderthals actually had no competitive advantage over each other [43]. The argument, based on a numerical model, is that, because AMH’s population was much larger than that of Neanderthals [44–46,12], AMH’s were statistically more likely to reach fixation in both Africa and Europe.

Although modelling is extensively used in studies of human evolution [e.g. 47–48], relatively few studies so far employed forward approaches on an explicit map to, for example, determine amounts of admixture, stepping stones or the most probably origin of AMH [49,50,38,51,52,43]. Kolodny and Feldman [43] used a simple “map” with only two demes to determine the chances that AMH would replace Neanderthals purely due to the larger population of AMH in Africa compared to Neanderthals in Europe. The SPLATCHE2 simulations [52] and the similar model by Eriksson et al. [38] assumed *a priori* that AMH have some advantage and explored the advance of AMH in relation to climate factors. These various models have in common that the species (AMH) and its competitive advantage are predefined.

Here we present a basic numerical model to simulate the spatial and temporal differentiation/speciation and the emergence, frequency and patterns of migration waves. Contrary to the above-mentioned models, no human species are defined *a priori*. The only aim is to explore the statistics of patterns, without claiming or attempting to simulate the actual emergence and spread of AMH, which would only be one of an infinite number of realisations of the stochastic model. Although a range of environmental factors have been invoked to explain various aspects of human evolution, we expressly do not include these in the models presented here. The aim is to provide a null hypothesis against which additional environmental factors and influences can be further tested.

## Methods

The model is based on a regular 2-dimensional grid of demes, each a square area with a number of individuals. The genome (G) of a deme is defined by a single string of *N* binary genes, with two alleles, either zero or one (similar to e.g. [53]. This genome is the single dominant or representative genome for the population of a deme. The temporal and spatial evolution of genomes is modelled in discrete time steps of length *Δt* in which mutations take place and genes are transferred between neighbouring demes. The distribution of genomes is visualised in RGB colour maps in which each deme is one pixel. Red is proportional to the number of ones in the deme’s genome, blue proportional to the number of genes identical to Gi={10101010, etc.} and green proportional to the number of genes identical to Gi={00110011, etc.}. Colours are stretched to maximise the colour range.

### Mutations

Mutations are carried out at the start of each time step. For each mutation, a deme is first randomly picked. Next, a random gene that has allele “0” everywhere in the model is chosen. This gene will be mutated by changing its allele from “0” to “1” in the deme. This procedure is repeated *M* times (the mutation rate) per time step. The mutation chance (*m*) per time step for one single deme is *m*=*M*/*A*, with *A* the area of the model. *m* is the chance that one mutation emanating from a single individual reaches fixation in the whole population (of size *N_D_*) of a single deme in a time step of *T* generations. *m* thus depends on *T, N_D_* (itself the product of population density *ρ* and area *S*^2^ of a deme) and the intrinsic mutation rate *m*_0_ of one individual per generation. The number of mutations that occur in one deme per generation is proportional to *N_D_·m*. However, the chance that a mutation reaches fixation within the population is roughly inversely proportional to *N_D_* for *s*≈0, with *s* the competitive advantage [54]. It follows that the rate of successful mutations within the deme is proportional to *m*_0_, and is not dependent on the population size of the deme. The model, however, uses time steps *T* of more than one generation. It will be shown below that *T*∝*N_D_*, and hence, *m*∝*m*_0_·*N_D_*.

We use the term “active mutations” for those mutations that have not reached fixation, i.e. they occupy at least one, but not every deme in the map of all demes Once a mutation has reached fixation by occupying every deme in the model it no longer plays an active role. The timing of fixation of a mutation is recorded, as well as the location of origin of the mutation.

### Allele exchange between neighbours

A mutation step is followed by one round of interbreeding in which single alleles may be transferred between neighbouring demes (here using a Von Neumann direct neighbourhood scheme). All demes are considered on average once in a random order for transfer between a deme (deme *a*) and a randomly selected direct neighbour (deme *b*). Each pair is thus treated every two steps on average. The chance that a transfer between neighbouring deme is considered in one time step is defined by the parameter *D*. It can be regarded as a general diffusion parameter: the chance, per time step, that an allele jumps from one deme to a neighbour in a random Brownian movement [55–57,54]. Assortative mating [58] is included in a way similar to the model of Barton and Riel-Salvatore [51]. With assortative mating, similar individuals are more likely to mate than different ones. This is implemented with the “assortative mating factor” (*α*), which reduces *D* as a function of Δ*G*, which is the number of genes that have different alleles in the two neighbouring demes under consideration:

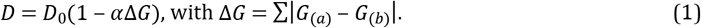

*D*_0_ is the reference diffusion parameter, equal to *D* for the case that *α*=0. It is first decided with a random-number generator whether a gene transfer will take place or not, according to Eq. (1). If this is to happen, all genes in the list are considered for allele transfer. The chance *P*(*a*→*b*) that an allele of the genome of deme *a* is copied to deme *b* is calculated with:

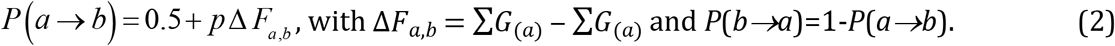

Δ*F_a,b_* is the difference in number of mutations (ones) between the two demes. The chance that a deme passes on its mutations to a neighbour is thus determined by the overall number of mutations relative to that neighbour (Δ*F_a,b_*) and the advantage factor *p* that defines the competitive advantage of these mutations (assumed the same for all mutations). In this paper we define the sum of all genes (*F*=Σ(*G*)) as the “fitness” of a deme. This is because the number of advantageous genes (*F*), multiplied by the advantage factor (*p*) determines the chance that genes are passed on to offspring. When *p*=0, mutations are neutral and there is no preferential transfer of alleles and we have purely random spreading of zeros and ones, but, on average, no change in their frequency. A positive value of *p* leads to fitter demes to pass on their alleles to their neighbours. Note that the transfer chance of a single allele does not only depend on its own value, but on that of the whole genome. This can potentially lead to less competitive zeros replacing more competitive ones.

To estimate the duration of one time step, we can consider the population of the two neighbouring demes as one during the transfer. Of interest is the case where the mutation only occurs in one of the two demes, i.e. when it is carried by 50% of the combined population. The time step *T* (in number of generations) is thus the time it takes for a mutation to reach fixation by genetic drift in the combined population with a chance of 0.5+*p* or extinction with a chance of 0.5-*p*. Using a basic Monte Carlo model for populations of up to a few hundred individuals, we find for the time step *T* (S1 Methods Eq. S3):

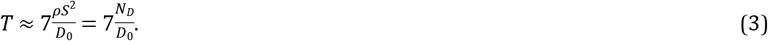

Here, *S* is the size of a deme and *ρ* the population density. Equation (3) only holds for weakly competitive mutations (*s*≪1), small values of p, and no variation in population density between demes. The factor *p* is related to *s*, the competitive advantage of the mutation [59,55–57,54] (S1 Method Eq. S2):

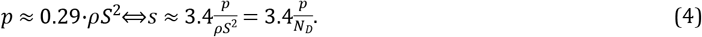

At a deme size of *S*=50 km and a population density of 0.01 individuals/km^2^, *T* is about 175 generations (≈4000 years at 25 years/generation) for *D*_0_=1 as used throughout this paper. Using *p*=0.05 results in a competitive advantage of *s*≈0.7%.

### Spreading of mutations

A single mutation in a field of demes without any other mutations has a chance of 0.5-*p* to disappear the first time an interbreeding event is considered with one of its neighbours. The initial survival rate is thus a function of *p*. The low initial survival rate is due to two factors. The first is a numerical effect that the width of diffusional front is less than can be resolved at the scale of the demes. The second is related to genetic drift [60–61], where the effective population (cluster of demes around the new mutation) is small and therefore the chance of survival smaller than when the mutation has spread over a large area.

Once a mutation has survived this initial nucleation phase by spreading over sufficient demes, the area occupied by the deme increases linearly with time. The expansion front is not sharp, but diffuse, in accordance with the diffusion-reaction model of [55]. The width of the diffusional front decreases with increasing *p* (Fig 1A-C) as the advantage factor becomes more important relative to the diffusional spreading. After the initial nucleation phase, the area (*A_mut_*) occupied by the single mutation increases linearly with the square of time. We define the velocity (*v*, in deme size per time step) of the expanding front as the rate of increase in radius (*r_eq_*) of an equivalent circle with area *A_mut_*:

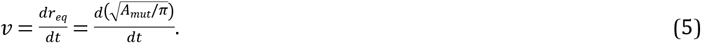

The spreading velocity (*v*_(*p*)_) as a function of *p* is determined from 2500 simulations of an expanding mutation after a stable diffusive front has been established (50<*r_eq_*<150 demes). *v*_(*p*)_ is determined by the chance (proportional to *p*) that the mutation is copied to a deme without that mutations and the length (*L_if_*) of the interface between demes with and without that mutation. *L_if_* is much larger than the circumference (2*πr_eq_*) of the equivalent circle in case of a diffusive front. *L_if_*/2*πr* thus decreases with *p* and can be approximated with a power law (using a least-squares best fit, with r^2^=0.99565; Fig 1D):

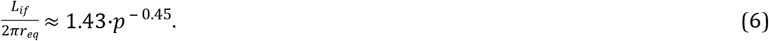

As a result, *v*_(*p*)_ is approximately a power-law function of *p* (r^2^=0.9993; Fig 1E):

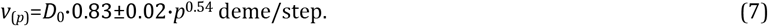

The exponent is slightly larger than 0.5 due to the fact that the spreading front becomes less fuzzy with increasing *p*. This means that demes at the front have fewer neighbours without the mutation when *p* is large than when it is small.

**Fig 1.**
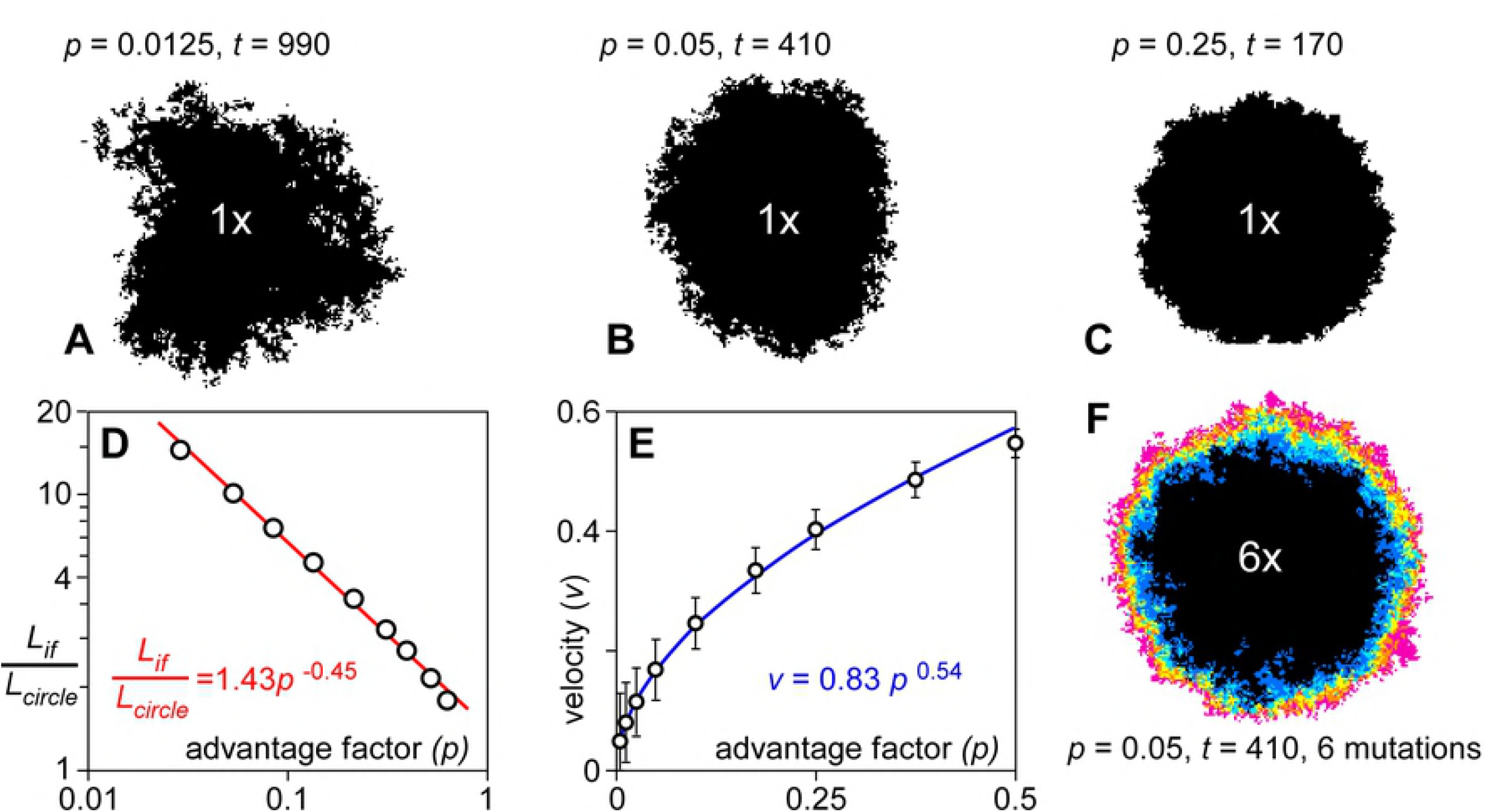
Spreading rates of mutations. Distribution of a single mutation originating from the centre of a circular model (*R*=100 demes) at the stage where the effective radius of the area occupied by the mutation is 60 demes. **A.** *p*=0.0125 at t=990. **B.** *p*=0.025 at *t*=720. **C.** *p*=0.05 at *t*=410. The effect of increasing *p* is an increase in spreading rate and a sharpening of the spreading front. **D.** Interface length divided by effective circle circumference versus drift factor in a double-log plot. Data approximately follow a power law. **E.** As a result, spreading velocity versus advantage factor also shows a power-law relationship. **F.** Distribution of six mutations seeded at the centre of the model (*R*=100 demes) at *t*=0. Colours show number of mutations on a deme from one (pink) to six (black).

The initial spreading velocity is higher when multiple mutations are placed in the centre of the model (Fig 1F). This is expected, because Δ*F*>1 in Eq. (2). However, the steady-state velocity of these mutations is the same when the diffusion front is wide enough (low *p*). The explanation is that demes within this diffusive front mostly only have a low Δ*F* with their direct neighbours and thus spread as fast as a single mutation. This implies that gene surfing due to high Δ*F* is not effective during steady-state spreading of an ensemble of mutations. However, when two different populations, with high Δ*F*, would suddenly come into contact the mutation spreading effect is expected to be high.

### Replacement events

Full genome transfers or replacement events are considered after the interbreeding step. Again, all demes and one random direct neighbour are considered in a random order. A full genome replacement is carried out if the absolute fitness difference |Δ*F*| is equal or greater than a set critical fitness difference Δ*F_crit_*. In that case, the full genome of the fittest deme is copied to that of its neighbour. After one round where all demes and one random neighbour are considered once for a replacement, the replacements may have led to new pairs that exceed Δ*F_crit_*. The routine is therefore repeated until Δ*F*<Δ*F_crit_* everywhere in the model. When Δ*t* is small, this semi-instantaneous spreading would be relatively fast (up to the order of a km/yr, depending on the size of demes). However, we use this scheme here to be able to track individual replacement “avalanches” within one time step. Contiguous areas that experienced a full genome replacement are termed “sweeps” here. The time and area of each sweep (*A_sw_*) is recorded at the end of each time step, as well as a map of all demes that experienced a genome replacement sweep.

### Aims

The aim of this paper is to illustrate the main types of evolutionary behaviour that result from different combinations of diffusive spreading with competitive advantageous mutations, assortative mating and replacement waves. It is not our intention to systematically investigate and quantify the effect of each of the parameters. We therefore only present four representative cases, using a reference model with 3 square inhabited areas, connected by narrow isthmuses. The areas or “continents” are 100×100, 50×50, and 25×25 demes in size. In each individual simulation presented below we keep all parameters constant in space and time. This implies that any environmental factors that could affect the competitive advantage of individual mutations, their spreading rate, and the population density are kept constant in space and time. This serves the aim of this paper to investigate patterns that develop in the complete absence of any environmental or other external influences. We consider this as a fundamental preliminary step in order to further discuss diversification processes during human evolution.

## Results and discussion

### Diffusion effect

We first consider the case (Fig 2A) without genome replacements (Δ*F_crit_*=∞) or assortative mating (*α*=0). The mutation rate *M* is set to four mutations per time step and *p* is 0.05. Individual mutations spread leading to increasing variation in genomes within the model. This is comparable to the geographic differentiation through isolation-by-distance as proposed for the reticulate or multiregional evolution model [62–63] or the recent assimilation model [64–65]. Demes in the centre of the model are on average nearer to the origins of mutations than are demes on the periphery. Fitness therefore increases faster in the centre than the periphery, as can be seen in the fitness profile across the three “continents” (Fig 2A). After steady outward fitness gradients are established at about t=250, genetic signatures migrate outwards, down the gradients. Signatures in the periphery are thus regularly overprinted by those coming from the centre.

**Fig 2.**
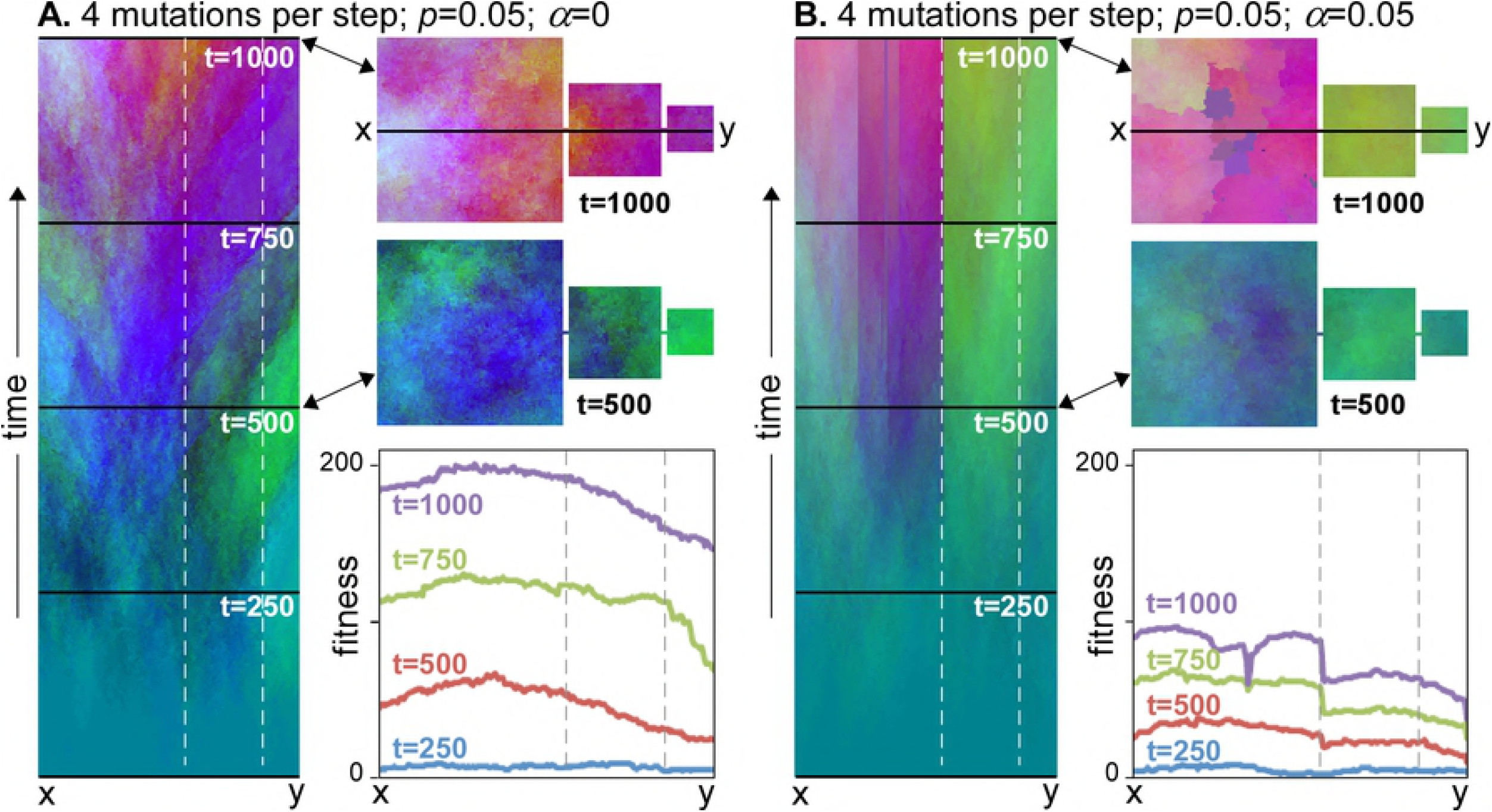
Time evolution over 1000 steps of genome signatures along a profile x-y through the three continents. Map view of genetic variations is shown at stages 500 and 1000. Colours qualitatively represent variations in genome, with the red tones proportional to “fitness”. Graphs show the fitness profiles for four time steps. **A.** In case of diffusion only, genetic signatures tend to emanate from the centre of the occupied areas and spread out towards the margins. **B.** When assortative mating is added, internally relative homogeneous “nation” regions with sharp boundaries develop from about *t*=500 steps. As these boundaries inhibit spreading of mutations, overall fitness increases more slowly than without assortative mating.

### The effect of assortative mating

The effect of the assortative mating factor (*α*) is to reduce the rate of gene exchange when the genomes of neighbouring demes are different. Figure 2B shows the effect of assortative mating in the same model as Fig 2A, but with *α* set to 0.05. This means decreasing genetic exchange up to Δ*G*=20, where exchange is reduced to zero. With initially low variations in genetic signature, the effect in the beginning is only to slow down differentiation. At about *t*=500, first neighbouring demes cease exchanging alleles, which allows their Δ*G* to increase further. This finally leads to homogeneous regions, bounded by fixed, sharp borders. In Fig. 2B these are visible as persistent, sharp changes in colour. New mutations cannot escape these “nations”. Because the fixation time within a “nation” is smaller than for the whole model, these mutations can now spread significantly before more mutations occur, thus keeping Δ*G* and Δ*F* low within a “nation”, while *F* for each “nation” keeps rising steadily. A large area, high mutation rate and low spreading rate (low *D*_0_ and *p*) all favour high values of both Δ*F* and Δ*G* (with Δ*G*≥Δ*F*). When these values remain too low, incipient borders shift and weaken again, which inhibits the establishment of permanent borders. This effect is visible in the medium and small “continents” that now behave as a closed system with highest fitness in the centre, but no internal “nation” borders. The development of “nations” or a structured population [66] results in a breakdown of the positive relationship (Fig 2A) between genetic signature and distance between points or isolation by distance [67]. This leads to a relative isolation of demes, which is strengthened through time.

### Effect of replacement sweeps

The effect of replacement is again illustrated with the three-continent model (Fig 3), using the same *p*=0.05 and *M* of four mutations per time step as before. Δ*F_crit_* is set at 10, so the genome of a deme is fully replaced by that of its neighbour if that neighbour has at least 10 mutations more.

**Fig 3.**
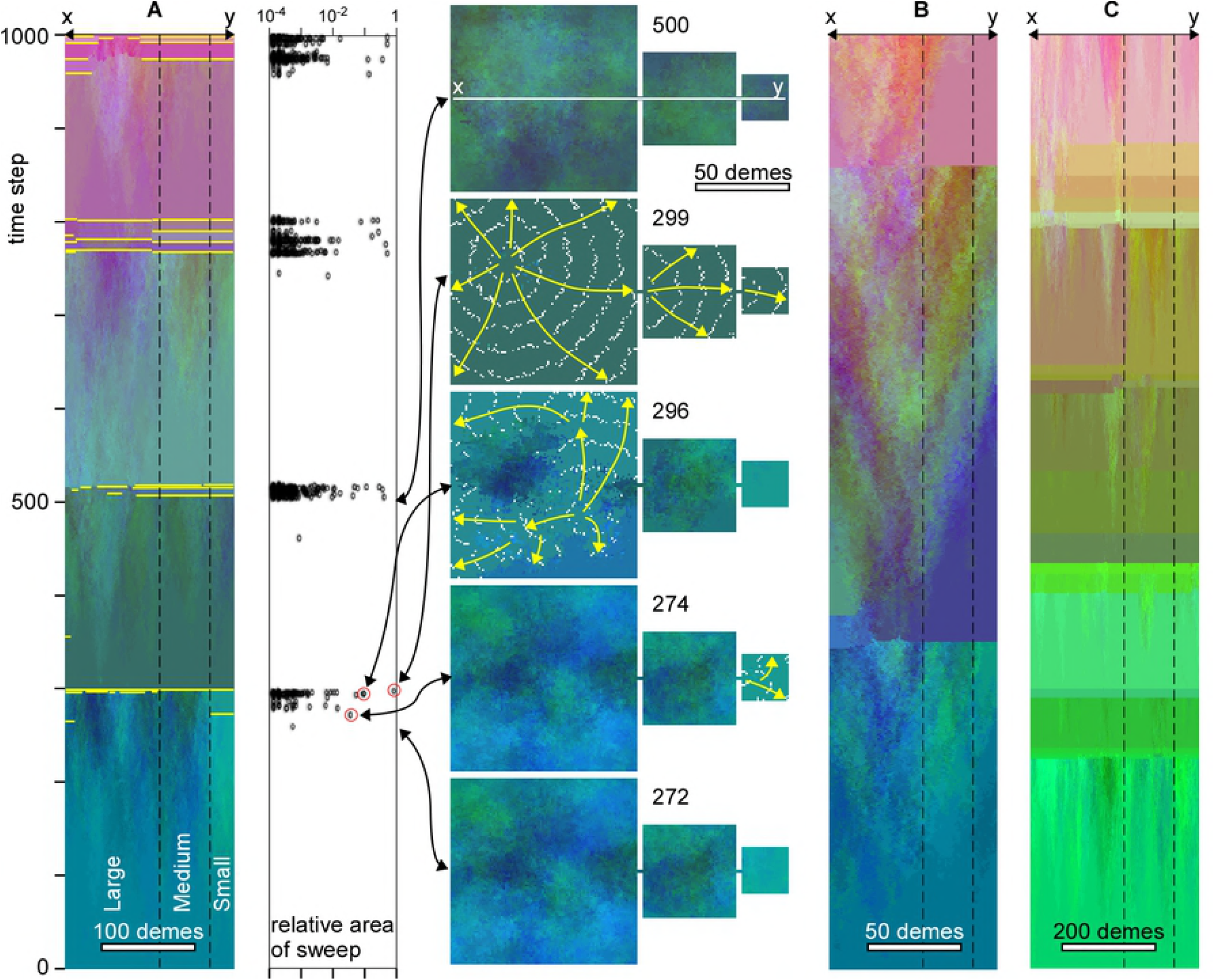
Evolution of genetic signatures in case of replacement sweeps in the absence of assortative mating. **A.** Temporal evolution along the line x-y through the centre of the model. Sharp changes in colour indicate replacement sweeps. The main ones are highlighted in yellow. Graph shows the area (relative to total area) of individual sweeps as a function of time. Replacement sweeps are strongly clustered in time. Map views show the genetic distribution at selected points in time. Direction and extent of selected sweeps are shown by yellow arrows and white lines that trace the front at regular intervals, respectively. Doubling (**B**) or halving (**C**) the deme size roughly doubles or halves, respectively, the average time interval between clusters of replacement sweeps, but not the general pattern.

In the absence of assortative mating (*α*=0), fitness increases steadily, especially in the centre of the model, until gradients exceeding Δ*F_crit_* are reached (at *t*=274 in Fig 3A). This typically happens somewhere between the centre and the margin. This is because, although fitness is highest in the centre, gradients are generally low here. Gradients are highest near the margins, where fitness is lower than in the centre. As a result, replacements first sweep the margins of the model (*t*=274), skirting the high-fitness centre (for example at *t*=296). Demes in the swept area with a single genome subsequently exchange alleles with the unswept demes, which rapidly leads to genomes with enhanced fitness again, and, hence, new sweeps (*t*=299). A rapid succession of admixture and replacement sweeps leads to homogenisation of the genome over the whole area. Although the new global genome is closest to that of the centre of the model, there are significant changes by admixture during the various successive sweeps.

After homogenisation of the genome, it takes some time for gradients to develop again to initiate a new cycle of replacement sweeps. This leads to a regular cycle of 200-300 time steps of differentiation without any sweeps, followed by a rapid succession of many small and a few large sweeps that sweep almost the whole model area (Fig. 3A). This pattern is in line with the punctuated-equilibrium model [68–70]. Reducing the resolution by a factor two, while keeping all other parameters the same, implies reducing the population density and the frequency of mutations in the model by a factor four. Gradients now increase at a lower rate and the duration of a full cycle is roughly doubled (Fig 3B). Doubling the resolution has the opposite effect and leads to a reduction of the cycle time (Fig 3C). Independent of resolution, admixture results in several large sweeps that together reset the genomes in the whole area.

Adding assortative mating (*α*=0.05) to the previous simulation significantly changes the evolution of the model (Fig 4A). After the initial differentiation period, first sweeps occur and, again, mostly sweep the margins. Contrary to the previous case where *α*=0, the sweeping genome is now unlikely to interbreed with fit demes at the edge of the swept area owing to their large Δ*G*. New sweeps are thus not immediately triggered for lack of admixture, and sweeps are less clustered in time. Demes in the centre are rarely or even never swept, providing a genetic continuity here. These demes have a higher fitness than the surrounding homogenised swept areas, and thus have a higher chance to initiate future sweeps. Marginal areas show distinct extinction events, as can be seen by distinct colour changes in Fig. 4B. Halving or doubling the resolution has the expected effect of increasing, respectively decreasing the time between sweeps (Fig 4B-C).

**Fig 4.**
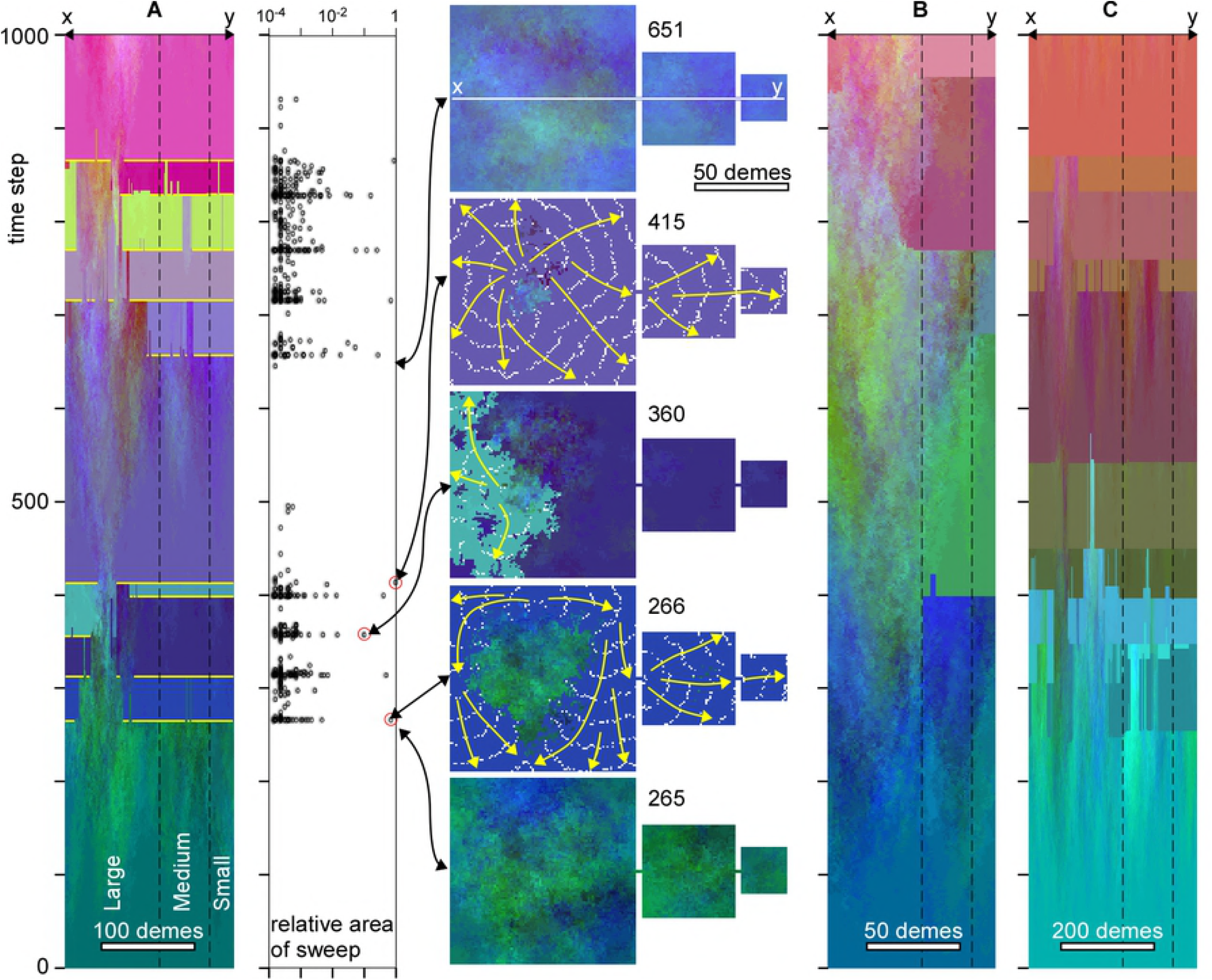
Evolution of genetic signatures in case of replacement sweeps as in Fig 3, but now with additional assortative mating. **A.** Temporal evolution along the line x-y through the centre of the model. Sharp changes in colour indicate replacement sweeps. The main ones are highlighted in yellow. Note the overall continuity in time of genomes in the centre of the large continent that is rarely swept by migration waves. Graph shows the area (relative to total area) of individual sweeps as a function of time. Replacement sweeps are less clustered in time than in case of no assortative mating. Map views show the genetic distribution at selected points in time. Direction and extent of selected sweeps are shown by yellow arrows and white lines that trace the front at regular intervals, respectively. Doubling (**B**) or halving (**C**) the deme size roughly doubles or halves, respectively, the average time interval between clusters of replacement sweeps, but not the general pattern.

Figures 3 and 4 show that marginal areas, in particular the small continent experience more sweeps than the centre of the area inside the large continent. The simulations shown in Figs 3A and 4A were also run for 10,000 steps, recording each time a deme was swept. Figure 5A shows that the chance for the two smaller continents and the margins of the large continent to be swept is about 1.5 times higher than in for the centre of the large continent in the absence of assortative mating. In case of assortative mating, the effect is even stronger. Demes in the centre of the largest continent thus have a much higher chance to be preserved, as these demes are rarely swept. This also affects the survival chance of mutations. The origins of mutations that reached fixation are plotted in Fig. 5B. We see that these are strongly concentrated in the centre of the large continent. While this continent occupies 76% of the whole model area, it is the origin of >99% of all mutations that reached fixation. The medium continent with 19% of the area delivered only <1% of all mutations that reached fixation and the small continent not a single one. The chance of a mutation from the medium continent to reach fixation is only 3.5% that of a mutation in the large continent in the simulation without assortative mating. In the simulation with assortative mating this change reduced to 1.7%. Replacement sweeps thus strongly favour the survival of mutations from the centre of the largest populated landmass.

**Fig 5.**
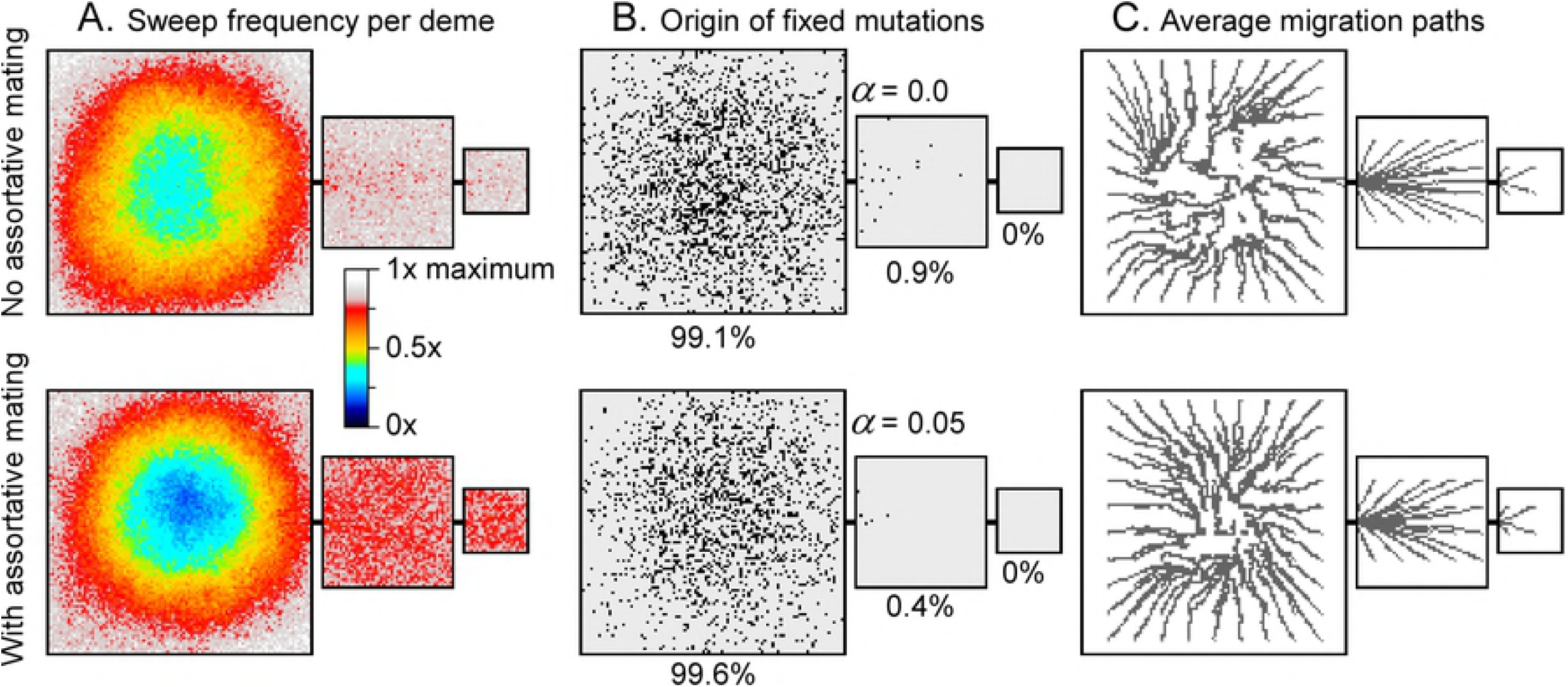
Origin of fixed mutations, as well as distribution and directions of sweeps. **A.** Origin of mutations that reached fixation shown at black dots on the map. Mutations that form in the middle of the large continent have a much larger chance of reaching fixation in the whole model area than mutations deriving from the margins, especially the small continent. **B.** Relative frequency that a deme is swept by a migration wave. Demes on the margins and small continents are swept more often than demes in the centre of the large continent. These patterns are more pronounced in case of assortative mating (below) relative to the run without assortative mating. **C.** Average directions of migrations within sweeps. Setting as in figures 3A (top) and 4A (bottom), run for 10,000 steps. C:

Directions of sweeps without (settings of Fig 3A) and with assortative mating (settings of Fig 4A) were recorded for simulations running 10,000 steps. Mean sweep propagation directions can be determined from this, and in turn, mean migration paths (Fig 5C). Migration paths consistently emanate from the centre of the large continent and lead to its margin and to the smaller continents. Migration directions are more consistent in case of assortative mating, resulting in a more consistent pattern of paths in the centre of the large continent.

### Fixation rate variation

After an initial period in which the system settles to a dynamic equilibrium, the number (*N_fix_*) of mutations that reach fixation (i.e. spreading over entire model area) increases linearly with time (Fig 6). In the absence of replacement sweeps, *N_fix_* increases steadily, whereas the *N_fix_*-time curve is stairway-like with replacement sweeps. This is because large replacement sweeps spread some mutations over large areas, resulting in a sudden, but temporary increase in fixation events. As the fitness landscape is flattened after these events, few mutations can subsequently reach fixation until the process is repeated again.

**Fig 6.**
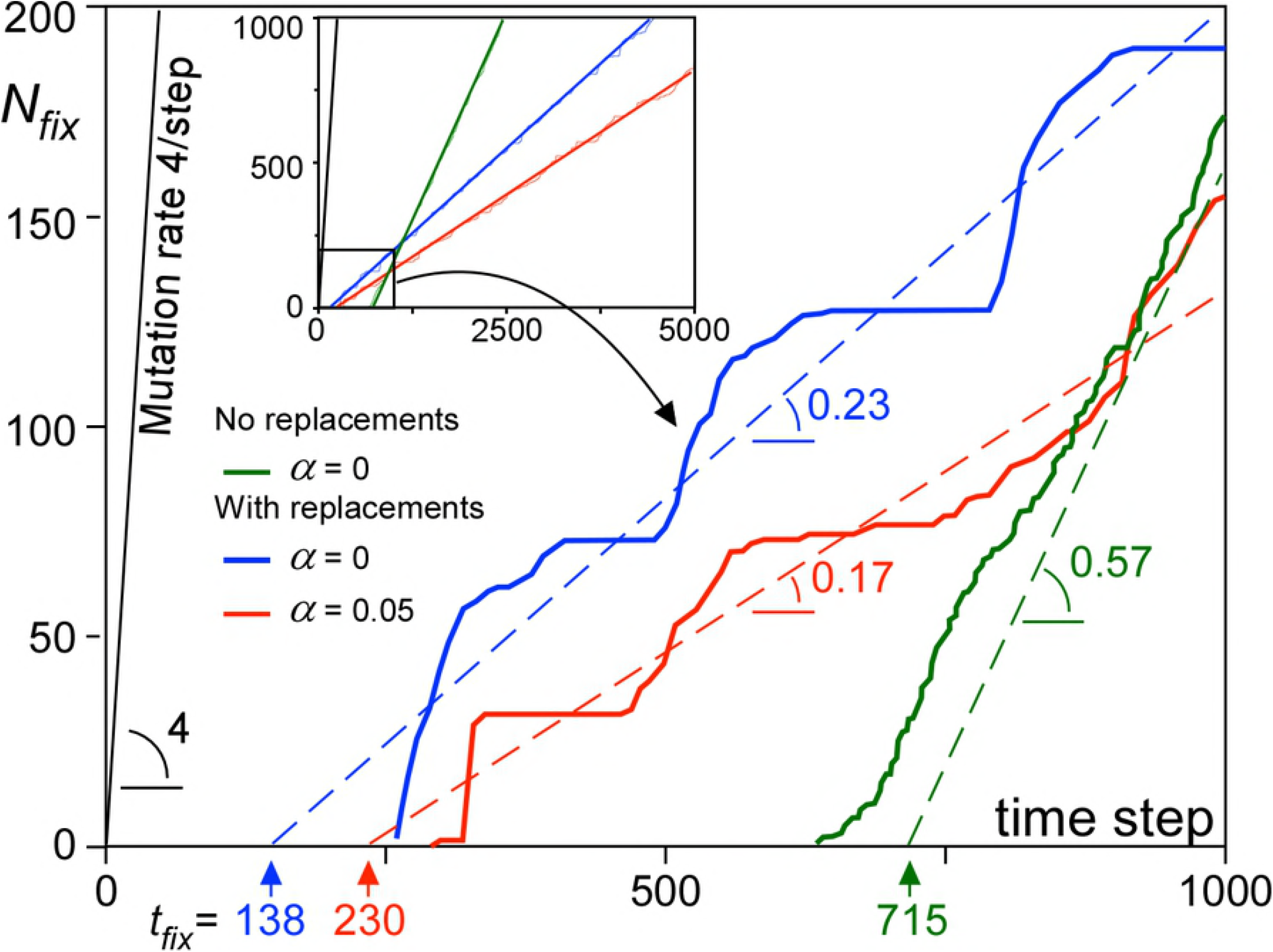
Number (*N_fix_*) of mutations that reached fixation as a function of time with linear regression lines (for *t*>600 steps). Main graph shows the first 1000 time steps, while the inset shows graphs for the full 5000 time steps on which the linear regressions are based. Intersection of the linear regressions is the mean time to fixation (*t_fix_*), while the slope is the rate at which mutations reach fixation. Replacement sweeps reduce the fixation chance of mutations by 60-70%, but also their *t_fix_* by 3-5 times.

The time-averaged fixation chance (*P_fix_*) of an individual mutation is derived from the slope of the *N_fix_*-time curve. *P_fix_* is significantly lowered by replacement sweeps. With interbreeding only, advantageous mutations are usually added to neighbouring genomes and few are lost by genetic drift before fixation, except at the nucleation phase just after the mutation occurred. Mutations in demes that are swept by replacement sweeps are lost, thus reducing the number of mutations that reach fixation [71]. Each wave causes a founder event, where only the limited genetic sample suddenly spreads over a large area [71–73]. The effect is more pronounced in case of assortative mating. The intersection of the linear regression of the *N_fix_*-time curve with the horizontal time axis (Fig 6) gives the mean time to fixation (*t_fix_*) for mutations that reach fixation. *t_fix_* is about 4 times smaller in case of replacement sweeps than without these. This is to be expected, as replacement sweeps provide an efficient means to spread mutations over the map.

The reduced *t_fix_* resulting from replacement sweeps has the advantage that a species can more quickly adapt to changes in the environment. This would cause some mutations to loose, and other to gain competitiveness. The latter can spread quickly in case of replacement sweeps. However, an inclination of a species towards replacements (low Δ*F_crit_*) comes at a cost. Replacement sweeps imply that part of the population is excluded and inhibited from further contributing to the species through their offspring. Furthermore, our simulations that are intentionally without any environmental changes show that a low *ΔF_crit_* also leads to spontaneous replacement sweeps in the complete absence of any external factors.

### Sweep area statistics

Although the largest sweeps are the most conspicuous in Figs 3 and 4, these are accompanied by many more smaller sweeps. Figure 7 shows the frequency (*f*) of sweep areas versus their normalised area (*A_n_* = sweep area/model area). We see that that the data plot on a straight line in the double-log plot, indicating a power-law relationship between *f* and *A_n_*, with an exponent -*q*:

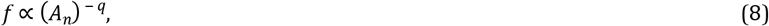

at least when *A_n_* is small (up to a few per cent of the total area). A power law (Eq. 8) was fitted to the data for *A_n_*<0.01 from simulations shown in Figs. 3 and 4, but run for up to 10,000 steps, resulting in *q*-values ranging from 1.81 to 2.09. Frequencies were normalised such that the power-law best fit frequency for *A_n_*=1 is unity in each of the six simulations. All these normalised data together overlap remarkably well (Fig 7). We obtain *q*=1.84 when applying a best fit to all data with *A_n_*<0.01. Normalised sweep areas >0.02 are overrepresented, with their frequencies up to >10x higher than the power-law trend. Notwithstanding this, small sweeps are orders of magnitude more common than the largest ones.

**Fig 7.**
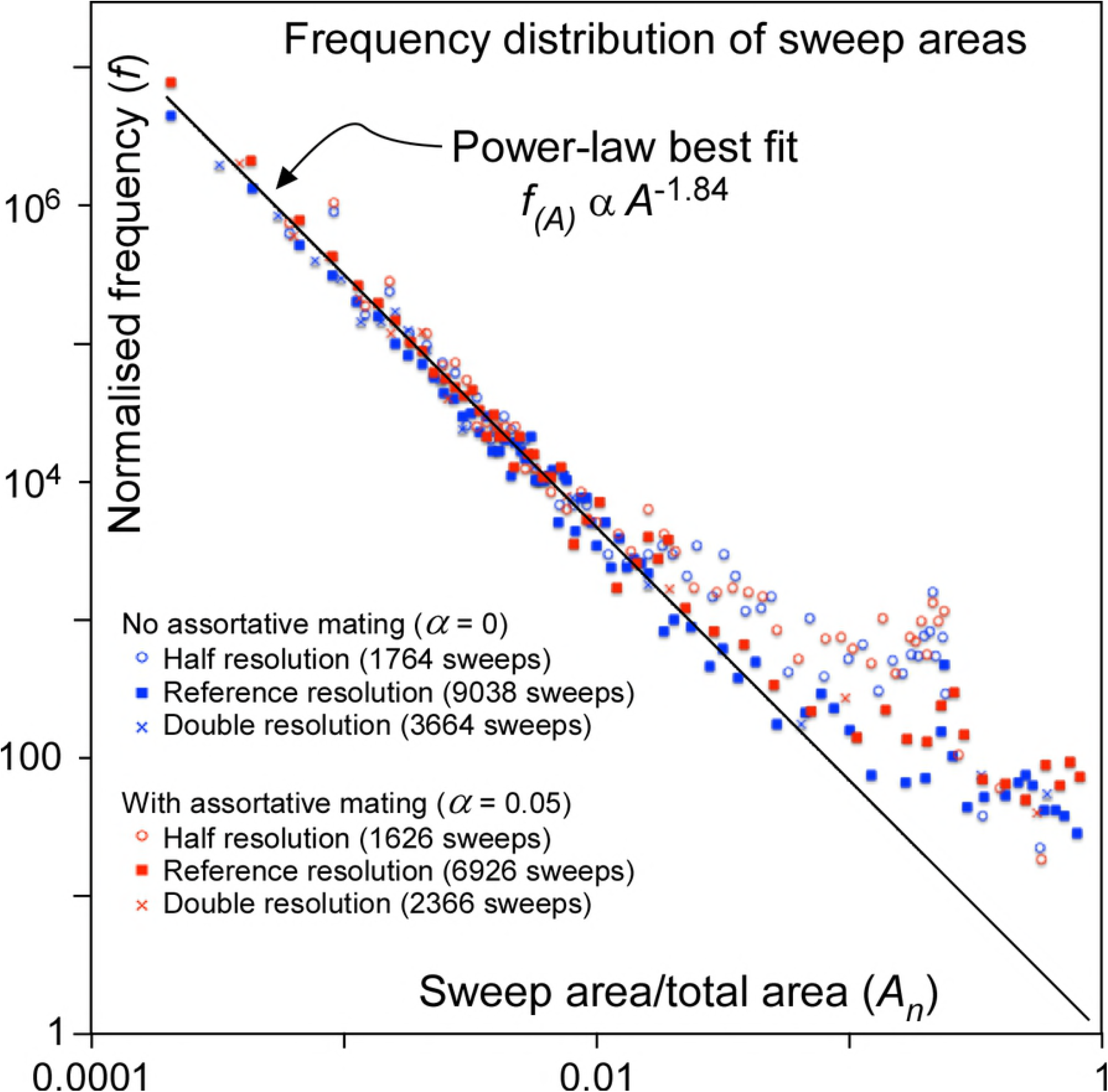
Normalised frequency distribution of areas swept by migration waves plotted against these areas (divided by total area) in a double-log plot. Data follow a power-law, except for the very largest sweeps. Dots represent the simulations with the schematic 3-continent map shown in figs. 3 and 4, but with 10,000 time steps.

The observed power-law relation is a hallmark of the self-organised criticality (SOC) model of Bak et al. [74] that has been used [75, 70] to explain punctuated equilibria [69]. The classical SOC model is the (numerical) sand pile in which grains are randomly sprinkled on a stage. Once critical heights of grain piles, or gradients in these heights are reached, grains are redistributed locally. One redistribution event can lead to neighbouring sites reaching the criterion for redistribution, sometimes leading to large “avalanches”. Sizes of these avalanches typically follow power-law distributions, as the critical state has no intrinsic time or length scale [74]. The current model is similar to the classical sand-pile model, as mutations are “sprinkled” on the map of demes. There is a criterion for redistribution (Δ*F_crit_*), which leads to replacement sweeps that indeed follow a power-law frequency distribution (Fig. 8). A first difference with the standard sand-pile model is that, contrarily to grains, mutations can multiply. The second is that the model includes diffusion as an additional transport mechanism for mutations. Models with two transport channels, one fast avalanche-like and one diffusional, have been applied to fluid flow through pores and fractures [76, 77], earthquake evolution [78, 79], and heat transport in plasmas [80]. These models show that such systems still exhibit SOC-characteristics, as long as the criterion for the fast transport is frequently reached. However, with increasing importance of diffusion, avalanches become more regularly spaced in time and larger, isolated events (so-called “dragon kings” [81, 82]) become more common [80]. This behaviour is indeed observed in our “mutation-pile” model, where large sweeps are over-represented compared to small ones and there is strong (*α*=0) and weak (*α*=0.05) cyclical behaviour with periods of semi-stasis (gradual diffusional differentiation), alternating with short periods of replacement sweeps. As such the model shows punctuated-equilibrium behaviour.

**Fig 8.**
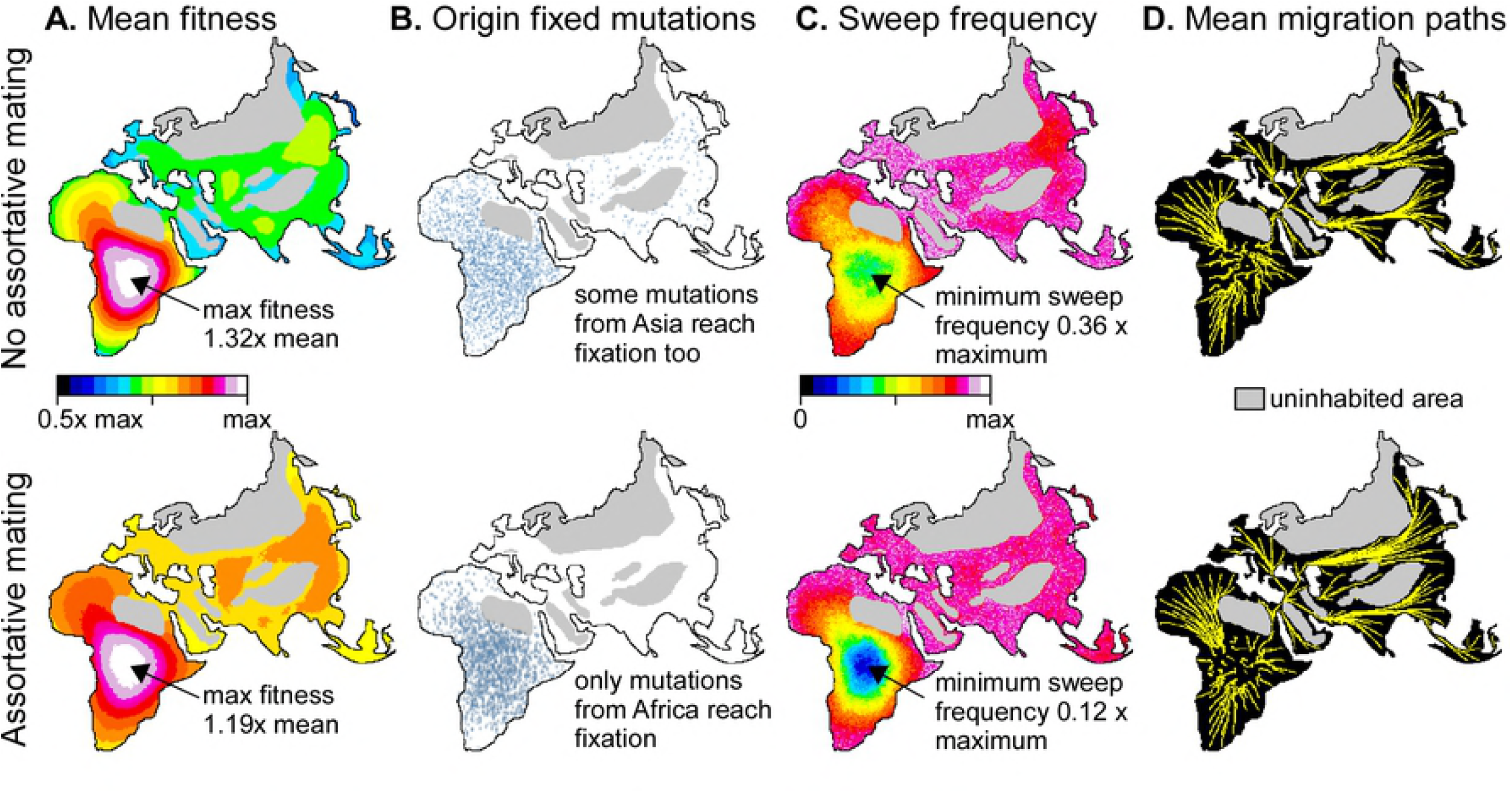
Example of applying the model for 10,000 steps with settings as in figures 3A and 4A to a map of the Old World. Top row without (a=0) and bottom row with assortative mating (*α*=0.05). **A.** Mean fitness of demes over the whole simulation, showing that demes in Central Africa are, on average, fitter than the minimum at the margins of the populated area. **B.** Origin of all mutations that reached fixation. Most (a =0) or all (a =0.05) these mutations originated in Africa. **C.** Number of times that a deme has been swept by a migration wave. Demes in Central Africa experienced significantly fewer sweeps than the rest of the world, especially in case of *α*=0.05. **D.** Mean migration paths, all emanating from central Africa and diverging towards the periphery of the continent and into Asia.

### Application to a world map

Having assessed the effects of, and patterns resulting from the different parameters on a schematic map with three continents, we now briefly illustrate the potential implication for human evolution. For this purpose we used a Fuller-projection map of the Old World (Fig 8), roughly adapted to ice age conditions by linking Japan and the British Isles to the mainland and assuming that large parts of northern Europe and Asia are (effectively) not inhabited. Although population densities (*ρ*) would never have been equal throughout the inhabited area, we maintain the assumption of a constant *ρ*, but excluded high-elevation areas (especially Tibet and the Pamirs) and desert areas in North Africa and the Arabian Peninsula, adapted from Eriksson et al. [38] for ~21 ka BP. In this configuration inhabited areas in Africa occupy 44% of the whole inhabited area. Inhabited areas evidently varied over time, but this simplified model serves the purpose of predicting the main expected patterns. Deme size was set at 50×50 km, resulting in 20808 populated demes, and settings were equal to those for the simulations shown in figures 3A and 4A. At a time step of about 4000 years (for *ρ*=0.01 individuals/km^2^), *p*=0.05 would lead to a mutation spreading rate of 0.002 km/yr (Eq. 5). Replacement sweeps in the current model take place within one time step. The maximum distance to travel, from South Africa to Japan is about 17,500 km, resulting in a maximum sweep velocity of ~4 km/yr. Such high velocities, however, only apply to the few very largest sweeps.

In both simulations, with and without assortative mating, mean fitness is highest in central Africa (Fig 8A). The fitness maximum is most distinct in case of *α*=0, because the long and regular interval between clusters of sweep events allow large-scale gradients to develop. Mutations that reach fixation mostly come from central Africa (Fig 8B). When *α*=0.05 no mutations that originated outside of Africa reach fixation. When *α*=0, a few mutations from Asia reach fixation, because sweeps from Africa can trigger “counter” sweeps after admixture with genomes from the margin of the swept areas. The chance that a deme in central Africa is swept by a migration event is higher in case of *α*=0 than when *α*=0.05 (Fig 8C). Despite these differences, migration directions are mostly emanating from central Africa in the direction of the margins of the occupied area (Fig 8D).

Sweep area frequencies follow the same power-law distribution (Eq. 8, Fig 9A) as in the abstract 3-continent model. Largest sweeps are again over-represented relative to the power-law trend for smaller sweeps. The effect becomes noticeable for sweeps that are larger than a few per cent of the total area. This is about 1/3 the area of Europe. The frequency distributions show two distinct peaks. One is at the area of Japan, which in the model forms a narrow peninsula connected to Asia. Sweeps apparently initiate at the connection with the peninsula (where mutations rarely occur) and then tend to sweep the whole peninsula. The second peak is at about 56% of the total area, which equals the area of Eurasia. This indicates that isthmuses, such as the Sinai Peninsula, play an important role and sweeps from Africa that entered Asia have an increased chance of then sweeping the whole of Eurasia.

**Fig 9.**
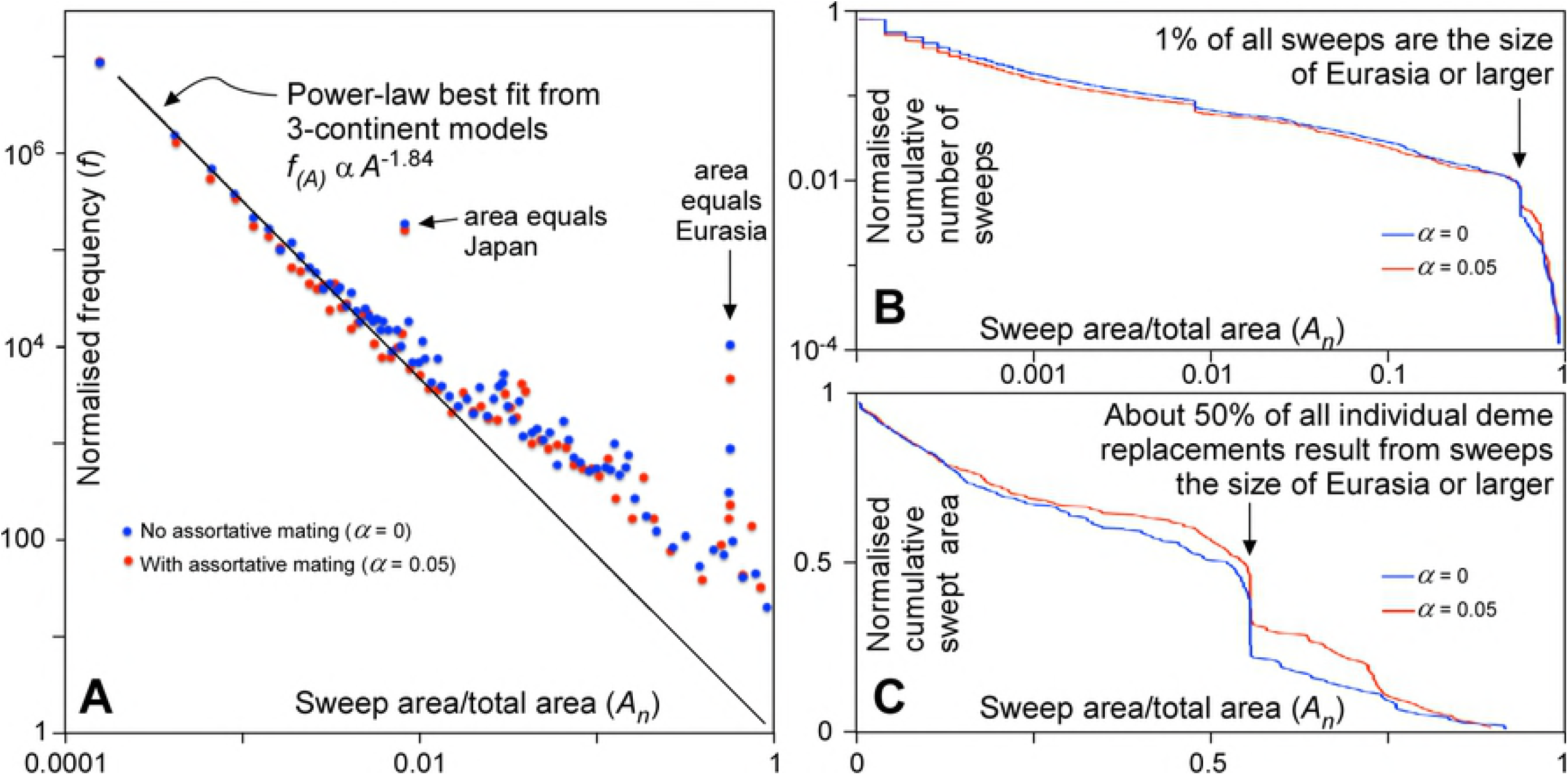
Sweep statistics of simulations for the Old World. **A.** Graph of normalised frequency as a function of sweep area. **B.** Cumulative graph of number of sweeps against sweep area. About 1% of all sweeps are the size of Eurasia (56% of total area) or larger. **C.** Cumulative graph of the areas swept as a function of sweep area. About 50% of all individual deme replacements result from sweeps are the size of Eurasia (56% of total area) or larger.

The cumulative number of sweeps as a function of area (Fig 9B) shows that sweeps of the size of Eurasia or larger represent about 1% of all sweeps. However, these few sweeps are responsible for about 50% of all individual deme replacement events over time (Fig 9C). If in the past there were one or two major out-of-Africa sweeps, one can deduce that there were in the order of 100-200 replacement sweeps in total. The average number of sweeps that an individual population in one single deme would have experienced would be about 2-4, double the number of very large sweeps. Two to four replacement events in a million years, i.e. ~40,000 generations, means that individuals in a deme have a chance in the order of 0.005-0.01% of experiencing a replacement sweep in their lifetime. Although sweeps thus rarely affect individuals, they have a profound effect on the evolution of the human genome, because about half these locally experienced sweeps are part global-scale sweeps that span a significant part of the whole populated Old World.

## Conclusions

Diffusional gene flow alone leads to a homogenisation of all populations [83]. However, ongoing mutations continuously produce local variations that take time to spread over the whole populated area. The combination of diffusion and mutations thus results in isolation by distance [67], with differences between local populations increasing with distance (Fig. 2A). The magnitude of these differences or clines depends on the balance between diffusion and mutation rate. The effect of assortative mating is to create a structured population [66] with relatively homogeneous “nation” regions, separated by distinct clines. While Scerri et al.[66] argue that the structuring is due to environmental and ecological drivers, our model (Fig 2B) shows that structuring can develop due to assortative mating without any such additional drivers. Without other evolutionary mechanisms, assortative mating reduces or inhibits exchange between regions and this exchange is restricted to neighbouring regions. Migration events are an efficient way to bring populations from far-removed regions into contact. Ensuing exchange across such new contacts leads to reticulate phylogenies [63].

Our model shows that, if population/species replacements do occur, replacement sweeps of all sizes up to the whole populated area are expected. The basic reason is that if one group of individuals can take over the area of their neighbours, the chance that they (or their offspring) can also take over the next area is larger than zero. This can, but must not, lead to “avalanches” of replacements that can span up to the whole populated area. Replacement-sweep-area frequencies systematically decrease as a power-law function of their area. For every world-spanning sweep, about two orders of magnitude smaller sweeps are to be expected, down to the size of one deme in the model. As expected, isthmuses appear to play a special “bottle neck” role during expansions. Once a sweep crosses an isthmus, there is an increased chance that the whole peninsula beyond is swept in its entirety. In simulations this leads to a distinct peak in the chance that the whole of Eurasia is swept in at an out-of-Africa event.

Replacement sweeps reduce the chance of fixation of mutations, but also significantly reduce the time to fixation of those mutations that do finally reach fixation. The propensity of a population to usurp the area of its neighbours if these are, for some reason, less competitive, has the benefit that advantageous mutations can quickly spread, for example after a change in environmental conditions. However, our simulations show that this propensity inevitably also leads to spontaneous replacement sweeps that are not triggered by any external factors but driven by genetic drift.

Our simulations show that replacement sweeps mostly emanate from the largest consolidated populated area. In the Old World this is Africa, especially during glaciation stages. The simulations indicate that the most likely origin for modern humans lies somewhere in (central) Africa, in line with what is deduced from the fossil record (e.g. [84]). However, East Asia also forms large and compact populated area, especially during warm periods, that would have been a second probable source for replacement sweeps. This emphasizes the need for further palaeoanthropological research in East Asia (e.g. [85, 5]).

Large migration sweeps generally emanate from the central regions of large compact areas and spread towards the margins. The spreading directions are mostly determined by coastlines, mountains and other uninhabited areas. This leads to a remarkable consistency of the directions, quite independent of the parameter settings (Fig 8C). The tendency to migrate towards coasts is consistent with the beachcomber model [86]. The spreading of a migration wave is more like that of an inkblot than along narrow, bifurcating migration paths that some authors envisage [87].

Mutations that arise in the centre of large compact areas have the highest change to survive and spread. This is not only because these areas are the probably sources of migration sweeps, but they are also the least likely to be swept themselves (Fig 8C). This pattern illustrates the relevance of the central areas as potential sources of new variants, but also in preserving old traits.

Considering a combination of semi-stasis periods alternating with replacement sweeps as a result of large-scale and many smaller expansions may contribute to understand the current biological distribution of human populations, as well as the variation in the hominin fossil record. For this reason we suggest an integration of concepts coming from punctuated equilibrium theory [68,70,75], multiregional postulates [24, 62, 63, 66], and current out-of-Africa migration models. Any migration wave that spanned the whole world is most likely to have come from Africa. If large migration waves can occur, more numerous smaller waves are to be expected too. Multiple out-of-Africa waves are, therefore, most likely if one happened. Although the propensity for replacement waves makes a species more adaptable to changes in its environment, a “side effect” is that such waves then also occur spontaneously. A spontaneous emergence and spread of modern humans from Africa should thus be regarded as a null hypothesis against which any hypothetical causes should be tested.

